# Predicting influenza H3N2 vaccine efficacy from evolution of the dominant epitope

**DOI:** 10.1101/300665

**Authors:** Melia E. Bonomo, Michael W. Deem

## Abstract

We predict vaccine efficacy with a measure of antigenic distance between influenza A(H3N2) and candidate vaccine viruses based on amino acid substitutions in the dominant epitopes. In 2016-2017, our model predicts 19% efficacy compared to 20% observed. This tool assists candidate vaccine selection by predicting human protection against circulating strains.

**40-word summary of main point:** Our *p*_epitope_ model predicts the ability of the influenza vaccine to reduce the A(H3N2) disease attack rate, with an r^2=0.77. This fast, sequence-based method compliments strain-to-strain antigenic comparisons from ferret models and provides antigenic comparisons for all circulating sequences.

## Text

Seasonal influenza constitutes a significant disease burden worldwide, with three to five million cases of severe illness and an estimated annual death toll of 290,000 to 650,000; however, vaccination can provide protection [1]. The vaccine component against influenza A(H3N2) is especially important, as increased morbidity and mortality are associated with this most commonly predominant subtype [2]. Influenza type A viruses are primarily recognized by the immune system via two proteins on their surface, hemagglutinin (HA) and neuraminidase [3]. These viruses constantly evolve to evade human antibodies, most notably by introducing amino acid substitutions into the HA binding sites, designated epitopes *A* through *E* in A(H3N2) (Supplementary Figure 1). Some epitopes appear to play a more dominant role during infection than others, and the one under the greatest immune pressure in a given season will have the highest percentage of amino acid substitutions, computed by dividing the number of substitutions by the total number of amino acids in that epitope [4]. Increased antigenic distance between the vaccine and infecting virus leads to decreased vaccine efficacy. Due to virus evolution, the World Health Organization (WHO) recommends the composition of the seasonal influenza vaccine twice a year based on the dominant influenza strains from the previous northern or southern hemisphere season [5]. Several antigenically similar candidate vaccine viruses (CVVs) and associated reassortants are made available. Generally, the reassorted viruses used in manufacturing the vaccine are antigenically identical to their CVV prototypes in the HA region [6]. For the 2016-2017 northern hemisphere influenza season, WHO recommended A/Hong Kong/4801/2014 (H3N2)-like CVVs [5]. However, vaccine efficacy for adults aged 18-64 against A(H3N2) influenza was only 20 ± 8% [2] (Supplementary Methods). Rather than an increased antigenic distance due to virus evolution, the 2016-2017 vaccine may have diverged from circulating viruses due to substitutions acquired during isolation of the CVV strains in eggs [7]. Passaging-related adaptations have posed an issue for A(H3N2) CVVs in particular [8,9] (Supplementary Table 3).

We have previously derived a statistical mechanics model that captures the dynamics of human antibody-mediated response to viral infection following vaccination [10]. In a recent application of our theory, we used the evolution of the dominant HA epitope to predict vaccine efficacy, the ability of the vaccine to reduce the disease attack rate [4]. Here we generalize the model to A(H3N2) data both from the 1971-1972 to 2015-2016 influenza seasons and from highly consistent, laboratory-confirmed studies over the past decade. Additionally, we expand the prediction to encompass average efficacy against an abundance of diverse A(H3N2) strains for a given season. We apply this novel approach to quantify antigenic distance and expected efficacy of the egg-adapted CVVs against all strains circulating during the 2016-2017 and early 2017-2018 seasons.

## Methods

The epitope-based dependence of vaccine efficacy in our model comes from a measure of antigenic distance termed *p*_epitope_, where

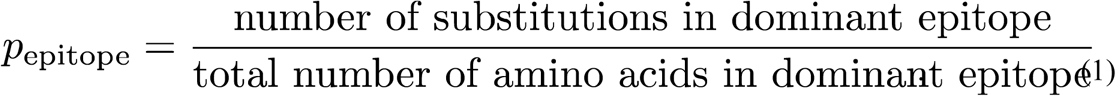

In ferret models, the typical measure of antigenic distance *d*_1_ is the log_2_ difference between vaccine antiserum titer against itself and the vaccine antiserum titer against a strain representative of the dominant circulating viruses [11]. A second measure d_2_ is the square-root ratio of the product of the homologous titers to the product of the heterologous titers [12]. We perform linear regression with vaccine efficacy data reported by WHO collaborating centers to compare the predictive power of *p*_epitope_ with these ferret-based distances. We then utilize our model to evaluate the primary protein structure of egg-adapted A/Hong Kong/4801/2014 CVV in the context of the 2016-2017 season [5] and to predict efficacy of the egg-adapted CVV for the newly recommended A/Singapore/INFIMH-16-0019/2016 (H3N2)-like vaccine [13]. Finally, we calculate the average efficacy of the CVVs against 6610 circulating A(H3N2) strains collected from September 2016 to November 2017, a unique capability of our *p*_epitope_ measure of antigenic distance in comparison to other clinical studies.

## Results

Our use of *p*_epitope_ to correlate antigenic distance with vaccine efficacy yields a coefficient of determination r^2^ = 0.77 on both data since 1971 and data over just the past 10 years (Figure 1). Conventional ferret *d*_1_ has r^2^ = 0.42 on data since 1971 and has dropped to r^2^ = 0.23 on data over the past 10 years. The extracted equation to predict vaccine efficacy *VE* is

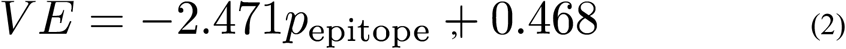

where the standard error on the slope is ±0.254 and the standard error on the y-intercept is ±0.032 (Supplementary Methods).

**Figure 1.**
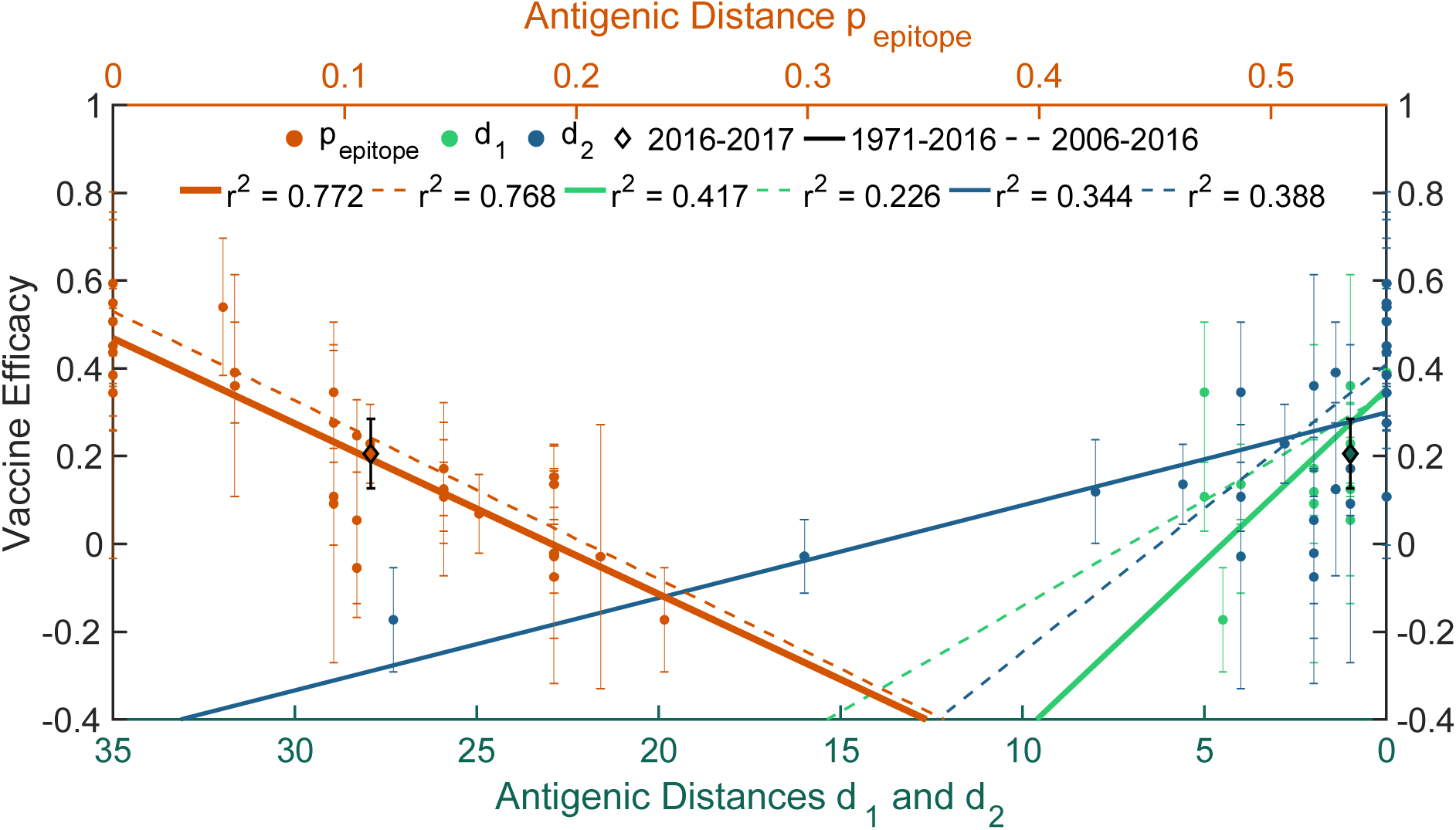
Influenza A(H3N2) vaccine efficacy as a function of three measures of antigenic distance. We plot *p*_epitope_ (red) from our model, d_1_ (green) and d_2_ (blue) from ferret serological studies [11,12], and the corresponding vaccine efficacy values for each influenza season. Linear regression of data from each measure of antigenic distance is shown for the past 45 years (solid lines) and for laboratory-confirmed studies in the past 10 years (dotted lines). The coefficients of determination (r^2^) are listed in the legend. Data for the 2016-2017 season are plotted separately (diamond points). The strong and consistent r^2^ between *p*_epitope_ and vaccine efficacy indicates that this model accurately predicts the data. We determined an average efficacy of 19 ± 4% for the 2016-2017 egg-adapted A/Hong Kong/4801/2014 vaccine, whereas 20 ± 8% was observed [2].

We calculate *p*_epitope_ = 0.095 between the egg-adapted A/Hong Kong CVV and A/Colorado/15/2014, a representative clade 3C.2a strain [7], and find the dominant epitope is *B*, with amino acid substitutions T160K and L194P. The reassorted CVVs contain some additional substitutions (Supplementary Figure 2), but as these substitutions are outside of the dominant epitope, we focus on the non-reassorted CVV for our analysis. Using Equation 2, we predict vaccine efficacy against this A/Colorado strain to be 23 ± 4% (Supplementary Table 1). The hemagglutination inhibition titers from post-infection ferret antisera reported for the 2016-2017 season [14] and our linear regressions of *d_1_* and *d_2_* predict vaccine efficacy to be 27 ± 5% and 28 ± 4%, respectively. Analysis of the reference A/Hong Kong strain, which was the clinical specimen collected from humans and expected to dominate in the population, reveals that this strain did not exhibit the epitope *B* substitutions. We predict it would have had a high 47 ± 3% efficacy against A/Colorado.

When comparing the CVVs to all circulating strains during the 2016-2017 and early 2017-2018 seasons, we considered both *p*_epitope_ (Equation 1) and the Hamming distance, which is the number of differing amino acids between each pair of strains in HA subunit 1 divided by the total 328 amino acids of the subunit. While the Hamming distance between the reference A/Hong Kong specimen and the consensus sequence of the circulating strains is 0.006, that of the egg-adapted CVV’s is 0.015 (Supplementary Figure 2). For the egg-adapted A/Hong Kong CVV, the average *p*_epitope_ is 0.111 due to substitutions in dominant epitope *B*, with an average predicted vaccine efficacy of 19 ± 4%. The reference A/Singapore/INFIMH-16-0019/2016 specimen has a Hamming distance of 0.003 to the consensus strain, smaller than the reference A/Hong Kong specimen has, and that of its egg-adapted CVV is 0.012. We calculate that the egg-passaged A/Singapore CVV has an average *p*_epitope_ of 0.118, leading to a predicted average vaccine efficacy of 18 ± 4%.

## Discussion

We quantify influenza virus evolution and predict the ability of the CVV to reduce the disease attack rate using our *p*_epitope_ measure of antigenic distance. While other factors influence real-world vaccine effectiveness in humans, *p*_epitope_ has accounted for much of the variance in CVV efficacy over the past 10 and 45 years (r^2^ = 0.77). The low vaccine performance against A(H3N2) during 2016-2017 appears to have been caused by the T160K and L194P substitutions that occurred in the dominant epitope *B* during egg-isolation of the A/Hong Kong CVV. Indeed, human antisera studies showed that individuals who had received the Flublok recombinant influenza vaccine (containing T160 and L194) that had been passaged in insect cells produced more effective antisera than did those who had received the inactivated influenza vaccines Fluzone or Flucelvax (containing K160 and P194) [7], which were both made from CVVs initially isolated in eggs [9]. Arguably, HA stability did not account for the larger response induced by Flublok, since all three vaccines induced a similar antisera response when challenged by K160-containing viruses [7].

It is critical to choose a vaccine strain with the minimal antigenic distance from all circulating A(H3N2) strains. While the reference A/Singapore specimen minimizes the Hamming distance, the reference A/Hong Kong specimen minimizes *p*_epitope_ (Supplementary Figure 3). WHO changed the recommended CVVs for the southern hemisphere because the egg-passaged A/Singapore CVV contains an N121K substitution that matches the majority of recent viruses, and ferret antisera were more successfully raised [13]. However, the egg-passaged A/Singapore CVV has two substitutions in residue *B* (T160K and L194P) that do not match the majority of circulating strains. We predict that this A/Singapore CVV will have comparable efficacy to the egg-passaged A/Hong Kong CVVs in humans.

The *p*_epitope_ of a pair of strains can be calculated nearly instantaneously or averaged for one strain against 6000 in seconds. By contrast, ferret models are restricted to a few analysis pairs, antisera production takes 3-5 weeks, and there is added difficultly due to strict biocontainment measures [9]. Additionally, *p*_epitope_ theory suggests which sequence discrepancies are contributing most to a lowered vaccine efficacy. We currently do not explicitly consider N-linked glycosylation, though *p*_epitope_ implicitly incorporates this seeing as discrepancies in glycosylation between two strains are generally due to amino acid substitutions, *e.g.*, T160K caused an observed lack of glycosylation site in the A/Hong Kong CVV as compared to circulating 3C.2a strains [7]. Our model can rapidly narrow down clinical specimens that are representative of viruses that will dominate the upcoming flu season. It can then be used in cooperation with ferret models to confirm that the CVVs and their associated reassortants have not acquired critical antigenic changes before the vaccine is manufactured and administered to the human population worldwide.

## Acknowledgments

Influenza strain sequences were retrieved from the open access EpiFlu database hosted by GISAID: Global initiative on sharing all influenza data, accessible at http://platform.gisaid.org/. See Supplementary Table 2 for specific contributions of both the submitting and the originating laboratories.

